# Endogenous flow and dialytaxis govern aging of Adenosine 5’-triphosphate (ATP) condensates

**DOI:** 10.1101/2025.08.23.671873

**Authors:** Feipeng Chen, Alexander J. Dear, Parrik Dang Kow, Yuchao Wang, Thomas C.T. Michaels, Ho Cheung Shum

**Affiliations:** Department of Mechanical Engineering, The University of Hong Kong, Pokfulam Road, Hong Kong (SAR), China; Advanced Biomedical Instrumentation Centre, Hong Kong Science Park, Shatin, New; Department of Biology, Institute of Biochemistry, ETH Zurich; Zurich, Switzerland; Bringing Materials to Life Initiative, ETH Zurich; Zurich, Switzerland; Department of Biomedical Engineering & Department of Chemistry, City University of Hong Kong, Kowloon, Hong Kong (SAR), China; Department of Chemical and Biological engineering, Northwestern University, Evanston, IL 60208, USA

## Abstract

Biomolecular condensates are increasingly recognized as pivotal regulators of cellular physiology, yet their pathological aging underlies numerous diseases. However, the mechanisms governing this condensate aging remain largely elusive. Liquid-liquid phase separation of minimal biomolecular building blocks, such as small peptides or nucleotides, offers ideal model systems to dissect the mechanisms underlying condensate formation and aging. Here, we report that condensates formed by Adenosine 5’-triphosphate (ATP) undergo liquid-to-solid phase transitions (LSPT) in macromolecularly crowded environments, evolving from dynamic liquid droplets into urchin-like fibrillar aggregates. In the initial stages of aging, small aggregates actively engulf surrounding droplets via direct contact and rapid wetting-driven merging. As aging progresses, internal flows emerge within condensates, with velocities oriented toward the nearest aggregate core. These flows arise endogenously from ATP fibrillization through the Marangoni effect and result in a long-range chemotaxis ripening process that facilitates transport of ATP molecules from peripheral droplets to a central aggregate. The Marangoni effect also drives the long-range motion of liquid droplets on hydrophobic surfaces towards the aggregates, representing a novel form of dialytaxis. These findings provide crucial and previously unrecognized dynamic behaviors and insights into the physical principles underlying condensate aging.

## Introduction

Biomolecular condensates function as dynamic hubs that spatiotemporally compartmentalize cellular spaces and regulate various biological functions, such as gene transcription [1, 2], synaptic transmission [3, 4], and noise buffering [5-7]. Formed via multivalent interactions between constituent molecules, these membrane-less condensates exhibit diverse physicochemical properties [8-11]. In healthy states, condensates typically exist as liquid droplets with dynamic microenvironments that selectively concentrate biomolecules and orchestrate countless enzymatic reactions [12, 13]. Under pathological conditions, however, condensates undergo dysfunctional transitions to form solid-like aggregates or gels [14, 15]. Such liquid-to-solid phase transitions (LSPT) or aging processes are increasingly linked to the pathogenesis of neurodegenerative diseases and cancers [16, 17]. Despite their critical roles in diseases, the molecular mechanisms governing the condensate aging remain poorly understood.

Previous studies have shown that droplet-like condensates formed by intrinsically disordered proteins (IDPs), such as FUS or hnRNPA1, can transition into glassy gels [15, 18-20]. These aging processes that unfold slowly over days have been described by the Maxwell model, suggesting that condensates maintain self-similar architectures, but their molecular interactions are strengthened during aging [18, 19]. However, for condensates undergoing rapid aging and dramatic morphological transformations into fibrillar or spiny structures within hours or even minutes, the underlying aging dynamics and mechanisms remain unclear [14, 21].

Recent studies have increasingly revealed that liquid–liquid phase separation (LLPS) is not limited to long-chain biomacromolecules such as proteins and nucleic acids. Rather, even short nucleotides and peptides have been found to undergo phase separation under suitable conditions [22]. This realization opens new avenues for forming model condensate systems using these minimal building blocks to dissect the physical principles underlying condensate behaviors, including aging. Moreover, phase separation of such simple molecules is highly relevant to origin-of-life research and the design of biomimetic protocells. A key advantage of using minimal building blocks lies in their conceptual clarity, which enables isolation and quantification of the physicochemical forces that drive complex condensate behaviors like liquid-to-solid phase transitions.

In this study, we leverage Adenosine 5’-triphosphate (ATP)—a minimal and biologically ubiquitous nucleotide—as a model system to study how crowding-induced phase separation evolves into fibrillar aging. We first discover that droplet condensates formed by ATP in crowded environments can transition into urchin-like fibrillar aggregates within minutes. We show that these aggregates grow through two main mechanisms: droplet wetting-induced engulfment and long-range chemotaxis ripening or dialytaxis. Theoretical analysis and experiments further reveal that these complex aging behaviors arise endogenously from ATP fibrillization through the Marangoni effect, with maximum velocities of the fields scaling linearly with the Marangoni number. These findings uncover a previously unrecognized repertoire of complex dynamics during condensate aging.

## Results and discussion

### Aging dynamics of ATP condensates

Adenosine triphosphate (ATP) molecules form droplet condensates in aqueous solutions containing polyethylene glycol (PEG) and ethanol (Fig. 1A). Thermodynamically, ATP condensates arise due to excluded volume effects of macromolecular crowders, driven by a segregative phase separation mechanism (Fig. 1B). Intriguingly, these ATP droplet condensates undergo a rapid liquid-to-solid phase transition (LSPT) within minutes, evolving from droplets into urchin-like fibrillar aggregates (Fig. 1C and movie S1). This phase transition is driven mainly by reduced ATP solubility in the presence of ethanol, which triggers the nucleation and crystallization of ATP molecules (Fig. 1B) [23]. As shown by a phase diagram with ATP concentration fixed at 100 mM, this aging transition occurs more rapidly under high PEG and ethanol concentrations (Fig. 1D, red symbols) but not at low concentrations of these constituents (Fig. 1B, orange symbols). Confocal microscopy also reveals that ATP aggregates exhibit as blooming flower-like morphologies, growing upward in three-dimensional space (Fig. 1E).

**Fig 1.**
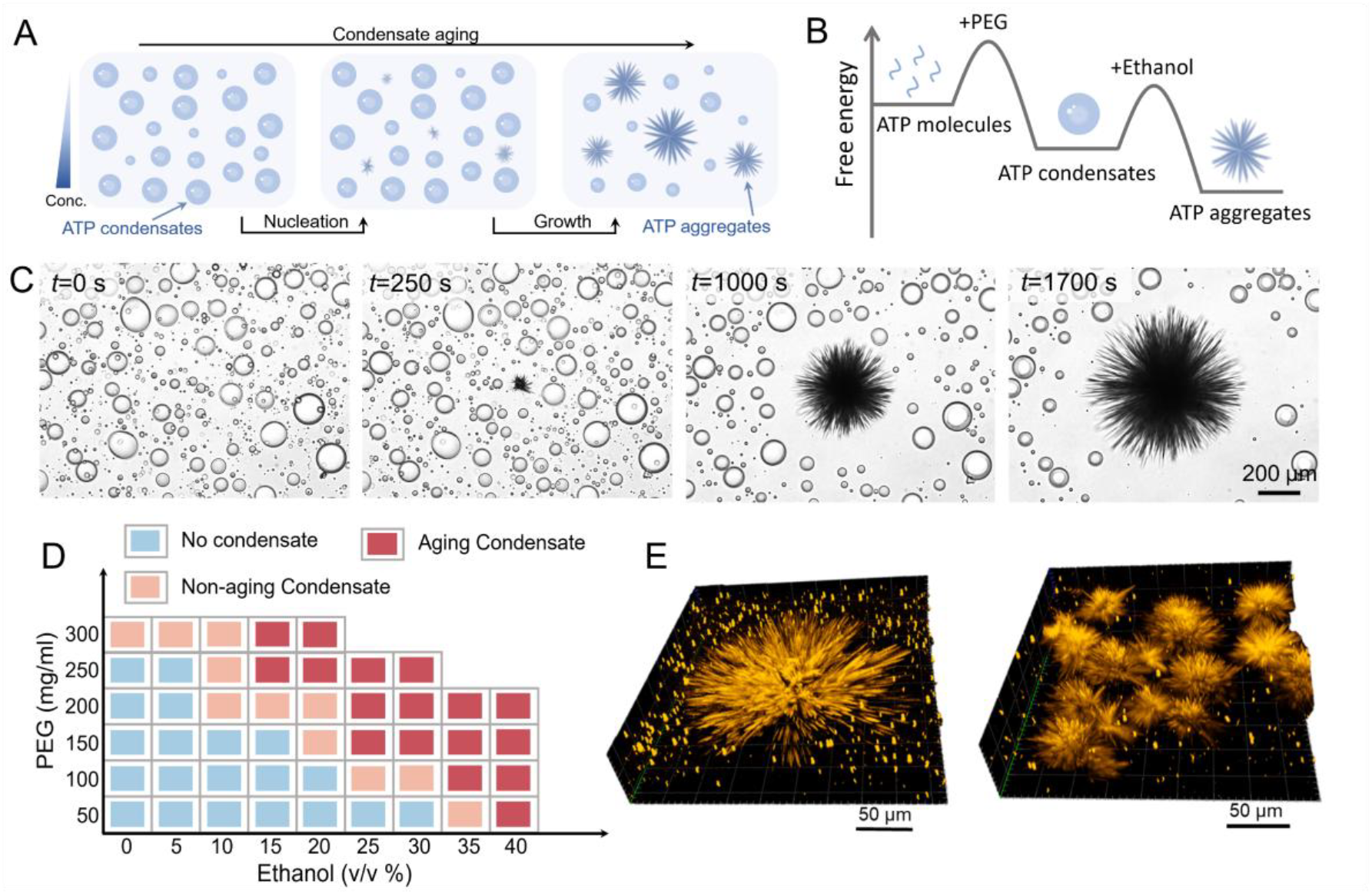
Aging dynamics of Adenosine triphosphate (ATP) condensates. (A) Schematic illustration of aging dynamics in ATP condensates. Some aggregates nucleate to continue growing during aging. (B) Schematic free energy landscape for the phase separation and aging of ATP condensates from kinetically trapped ATP monomers to intermediate state of droplet condensates, and then the thermodynamically stable ATP aggregates. (C) Bright-field images of an aging condensate over time (bottom: *c*_ATP_ = 100 mM, *c*_PEG_ = 250 mg/ml, and *c*_eth_ = 20 % v/v). (D) Phase diagram of ATP condensates as functions of PEG concentration (mg/ml) and ethanol volume fraction (v/v %), with ATP concentration fixed at 100 mM. Three regimes are denoted in the phase diagram: no condensate (blue squares), non-aging condensates (orange squares), and aging condensates (red squares). (E) 3D confocal fluorescence microscopy images of ATP aggregates exhibiting as blooming flower-like morphologies (labelled with Rhodamine-PEG).

During nucleation, a nascent aggregate of 50-80 µm in diameter forms in solution (Fig. 1C and movie S1). The aggregate growth involves two distinct phases: initially, it directly engulfs adjacent droplets upon contact, and later, it grows by depleting ATP molecules from distant droplets in a sink-like manner (Fig. 1C and movie S1). The later phase is reminiscent of Ostwald ripening in water-in-oil emulsions driven by molecular flux from smaller droplets to larger droplets driven by pressure gradients [24]. Notably, the entire aging transition proceeds orders of magnitude faster than previously reported condensate aging processes that typically unfold over days [14, 18, 21]. These results highlight new mechanisms driving the aging of ATP condensates.

We further interrogate how environmental conditions influence the aging dynamics of ATP condensates. Consistent with the phase diagram (Fig. 1D), aging is accelerated at higher ethanol and PEG concentrations (Fig. S1 and Fig. S2). However, high temperature suppresses aging presumably due to enhanced mobility and solubility of ATP molecules (Fig. S3 and Fig. S4). In addition, salt ions (NaCl, KCl, MgCl_2_, and CaCl_2_), 1,6-hexanediol (HD), and urea promote aging, as shown by increased numbers of ATP aggregates (Fig. S5-Fig. S10). Mechanistically, monovalent ions, 1,6-HD, and urea could reduce the release of counterions from ATP molecules, thereby promoting the neutralization and aggregation of ATP molecules [25]. Differently, divalent ions (e.g., Mg^2+^, Ca^2+^) could bridge ATP molecules to enhance their aggregation [26]. For consistency, we use a representative composition (*c*_ATP_ = 100 mM, *c*_PEG_ = 200 mg/ml, and *c*_eth_ = 25 % v/v) to study the aging dynamics of ATP condensates without additional additives in the following sections.

### Wetting-induced leaps of droplets and aggregates

After nucleation, aggregates extend needle-like fibrils that touch and engulf neighboring droplets (Fig. 2A). Strikingly, the wetting of aggregates by droplets induces rapid leaps and co-localization of droplets with aggregates at velocities of up to 10 µm/s (Fig. 2B, Fig. S11, and movie S2-S3). In addition, these aggregates also migrate along trajectories radiating from their centers to adjacent droplets and twist their bodies as they shuttle between droplets (Fig. 2B). To understand these wetting-induced leaps, we analyze forces generated during droplet-aggregate interactions (Fig. S11 and Movie S3). Assuming that droplets are approximately spherical with radius *R*, the drag force *F*_d_ on the droplet can be estimated using Stokes’ law as *F*_d_ = 6*π*μ*R*v, where μis the fluid viscosity and v is the translational velocity. The lack of inertial effects in the low-Reynolds number regime means that the drag force on the moving object at a steady state must equal the wetting force *F*_w_, i.e. *F*_d_ + *F*_w_ = 0. We estimate the drag force using a wetting event occurring at 02:14-02:16 in the movie S3. The droplet with radius *R* ≃ 12.5 µm moves approximately 10 µm over 1 second, yielding a velocity of approximately v ≃ 10 μm/*s*. This gives a wetting force of *F*_w_ ≃ 2.5 pN.

**Fig 2.**
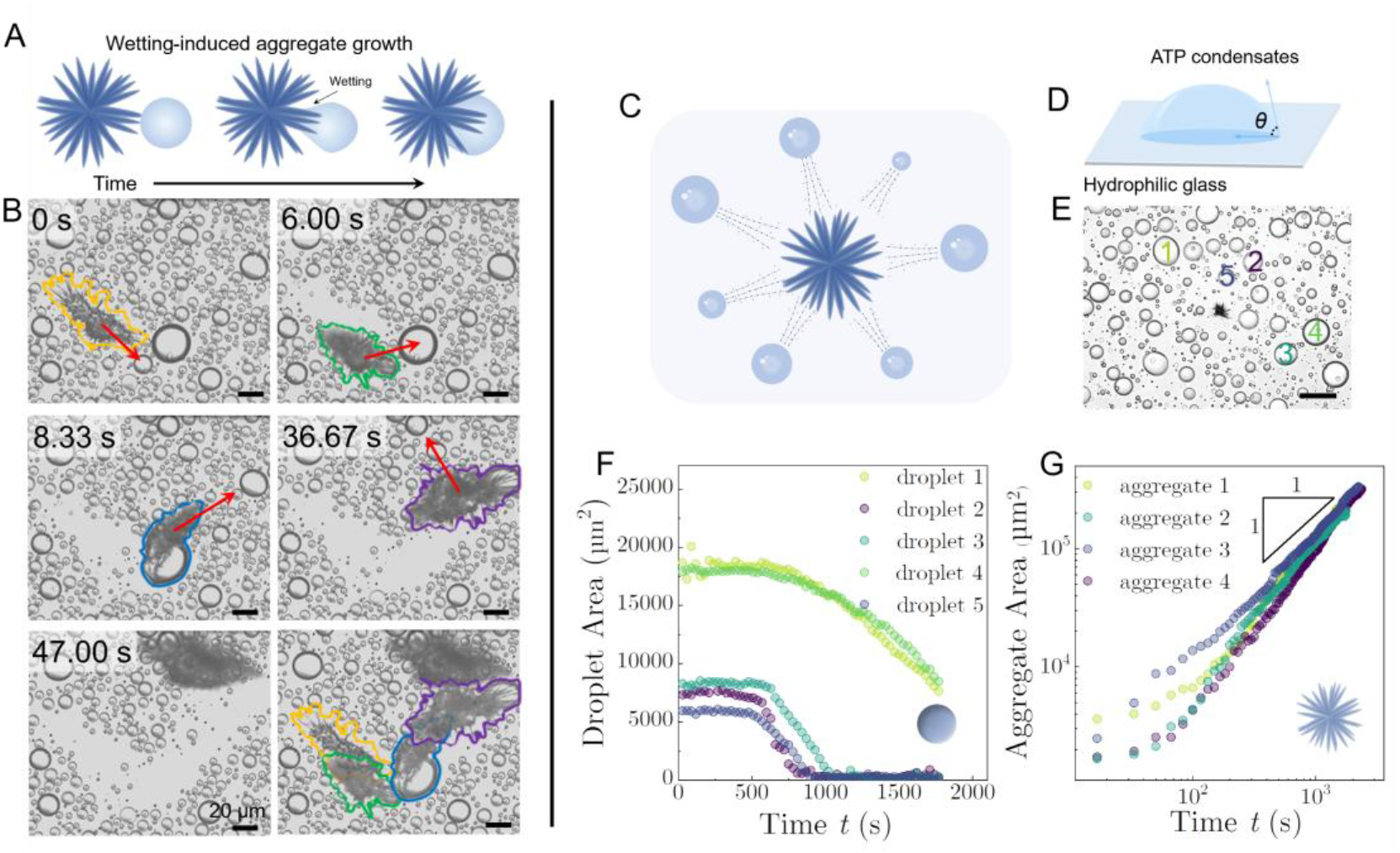
Wetting-induced leaps and long-range chemotaxis ripening during condensate aging. (A) Schematic illustration of wetting-induced leaps and colocalization of an aggregate with a droplet. (B) Bright-field images showing an aggregate leaping to sweep and engulf distant droplets over a long trajectory. The last image is a composite image showing the time-lapse positions of aggregates marked in different colors. (C) Schematic of long-range chemotaxis ripening in late-stage aging, enabling the transport of ATP molecules from neighboring droplets to a center aggregate. (D) Schematic illustration showing pinned condensates on hydrophilic surfaces. (E) Bright field image showing the positions and label numbers of droplet condensates surrounding a center aggregate condensate. The scale bar is 200 µm. (F)(G) Time-evolution of droplet and aggregate areas during the aging process.

We estimate the fibril thickness as approximately 0.5 µm. For fibrils with square cross-section profiles, this would correspond to a circumference *C* of 2 μm. The wetting force exerted by such fibril on droplets along the fibril length is on the order of *F*_w_ ∼ δ*γ* · *C*, where δ*γ* is the reduction in overall surface tension due to wetting. Using our calculated *F*_w_, this implies δ*γ* ∼ 10^−6^ N/m. Repeat calculations for multiple events in the Movie S3 show comparable δ*γ* values within the same range of 10^−6^ N/m. This estimated value is consistent with typical measured values of condensate surface tension of 10^−5^ − 10^−7^ N/m, validating our calculations and implying a typical wetting force in the pN range [27, 28]. These forces are sufficient to drive droplet leaps and their merging with aggregates.

### Long-range chemotaxis ripening of condensates

Once the aggregate has consumed all droplets in its immediate vicinity, its growth continues via a long-range chemotaxis ripening mechanism (Fig. 2C). On the hydrophilic glass substrate, all ATP condensates wet and remain pinned to the surface (Fig. 2D). As a result, the central aggregate continues to grow, while distant droplets shrink, indicating that ATP molecules diffuse from radially distributed droplets toward the aggregate (Fig. 2E and movie S1). To characterize the ripening kinetics, we track the projected area of droplets and aggregates over time (Fig. S12). Quantitative analysis yields clear trends: the areas of adjacent droplets decrease over time (Fig. 2F), while those of the central aggregates increase linearly with time (Fig. 2G), meaning that their characteristic radius *r* increases as *r*∼*t*^1/2^. Interestingly, this square-root time dependence of the aggregate size is explained by a diffusion-limited model, where the aggregate behaves as an absorbing sphere, with monomers integrating into its structure immediately upon contact (details in the supplementary materials). These aggregates thereby act as a sink, generating chemical gradients that drive long-range chemotaxis ripening of droplet condensates. Notably, diffusion-limited fibril growth is rarely seen in systems without liquid-liquid phase separation, e.g., amyloid fibers [29]. Therefore, the presence of liquid condensates causes a profound change in the kinetics of fibril growth.

### Directed flows inside aging ATP condensates towards the central aggregate

To visualize molecular diffusion from peripheral droplets to the central aggregate, we add fluorescent tracer microparticles into the system. These microparticles preferentially partition into droplet condensates (Fig. 3A). Notably, particle image velocimetry (PIV) analysis of tracer particles reveals the emergence of strong internal flows within aging condensates (Fig. 3A and movie S4), exhibiting counter-rotating vortices at the forefront of droplets with maximum flow velocities reaching ∼10 µm/s (Fig. 3A). Aging condensates there refer to those either actively transforming into fibrillar aggregates or situated in proximity to aggregates. In contrast, non-aging condensates, formed at lower ethanol concentration (20 % v/v), lack such internal flows (Fig. S13 and movie S5). In addition, flow velocities within transforming aging condensates fluctuate periodically, but their magnitudes decay over time as they mature into aggregates (Fig. 3B, red symbols and fig. S14). In comparison, non-aging condensates maintain negligible velocities throughout the observation period (Fig. 3B, blue symbols). Notably, in contrast to non-equilibrium condensate systems maintained by continuous chemical reactions [30, 31], our condensates systems are globally at equilibrium, while their internal flows arise endogenously from the fibrillization process of ATP molecules.

**Fig 3.**
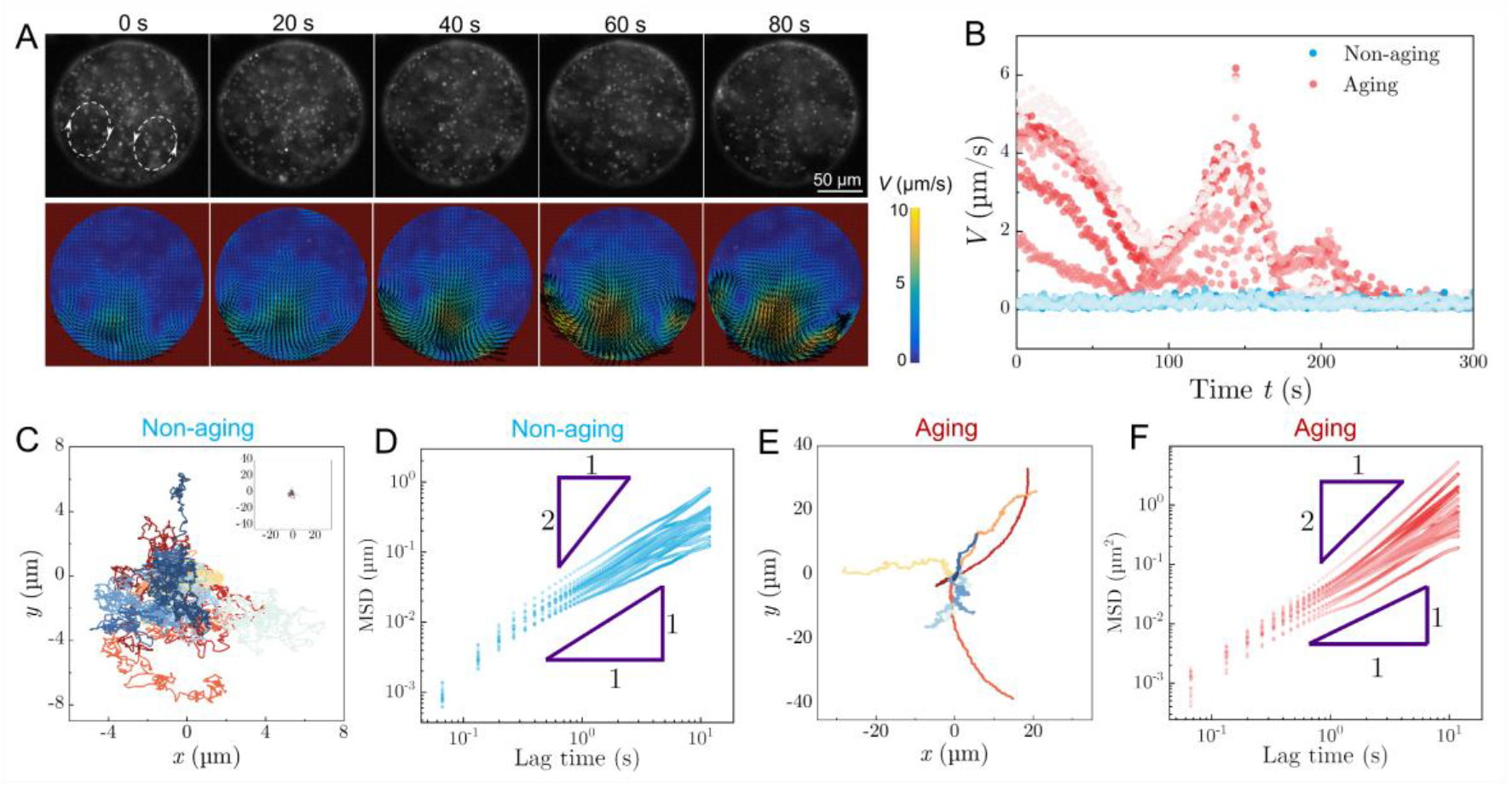
Active flows within aging droplet condensates. (A) Fluorescence images and particle image velocimetry (PIV) images of a droplet condensate close to an aggregate. Black arrows in the PIV images denote the velocity magnitude and direction of flow fields. (B) Time-evolution of flow velocities in aging condensate (red circles) and non-aging condensates (blue circles), with aging condensates undergoing complete droplet-to-aggregate transformation during measurement. (C) 2D trajectories of nanoparticles in non-aging condensates. (D) Mean squared displacement (MSD) as a function of lag time Δ*t* for nanoparticles moving inside non-aging condensates. in the non-aging condensate and aging condensates. (E) 2D trajectories of ballistic motions of nanoparticles in aging condensates. (F) Mean squared displacement (MSD) versus lag time Δ*t* for nanoparticles moving inside aging condensates. Different colors in each plot represent independent tracked particles.

Single-particle trajectory analysis further distinguishes these different dynamics. For example, microparticles in non-aging condensates meander randomly within local microdomains (Fig. 3C). By calculating the mean squared displacement (MSD), described by a power-law scaling as MSD(Δ*t*) = 4*D*Δ*t*^α^, where *α* is the scaling exponent and *D* is the diffusion coefficient, microparticles inside non-aging condensates exhibit classic Brownian motions with *α* ≈ 1 (Fig. 3D). In comparison, particles in aging condensates exhibit superdiffusive ballistic motions over long distances (Fig. 3E), with MSD exponent close to 2 (Fig. 3F). This quantitative shift in scaling, from *α* ≈ 1 (Brownian) to *α* ≈ 2 (ballistic), directly captures the transition from passive, unguided movements in non-aging condensates to active, directed transports in aging ones.

To understand how active flows drive long-range chemotaxis ripening, we analyze collective flow fields within neighboring droplet condensates surrounding an aggregate (Fig. 4A and Fig. 4B). It is observed that all neighboring droplets exhibit internal flows (Fig. 4C and movie S6-S7). In particular, velocity vectors *θ*_v_ at the forefront of these droplet condensates uniformly orient toward the central aggregate (Fig. 4D). By comparing the angular difference between each droplet’s instantaneous velocity *θ*_v_ and its azimuth angle relative to the aggregate *θ*_d_, it shows the distribution of *θ*_v_-*θ*_d_ is centered at -180° (Fig. 4E), confirming nearly perfect radial inward flow. We further quantify the relationship between their flow strength and spatial positions. Plotting the maximum internal velocity *V*_*c*_ of each droplet against its distance from the aggregate *d* reveals a general decrease in *V*_*c*_ over *d*, though data from three independent experiments remain scattered (Fig. 4F).

**Fig 4.**
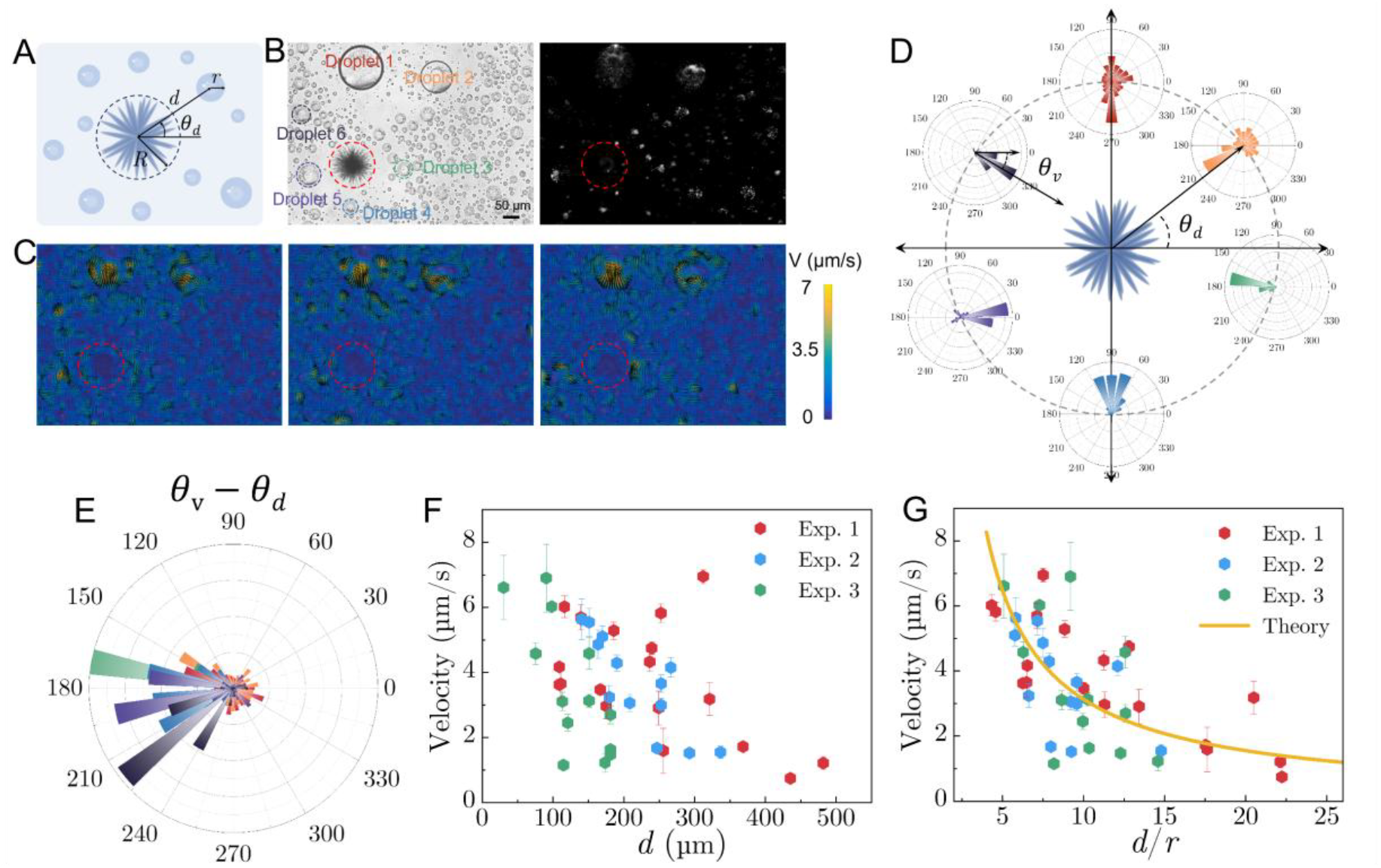
Collective active flows in neighboring droplets toward a central aggregate. (A) Schematic of the setup for Marangoni flow analysis: a spherical droplet of radius of *r* is located at a center-to-center distance *d* and an azimuth angle of *θ*_d_ from the fibril aggregate of radius *R*. (B) Bright-field and fluorescence images showing six neighboring droplets surrounding a central aggregate marked by red dashed circles. (C) PIV image of flow fields within the solution. Black arrows denote the velocity magnitude and direction of flow fields. (D) Polar histograms of velocity angles for six droplet condensates positioned at different azimuth angles relative to the aggregate. (E) Polar histograms of angular difference *θ*_v_-*θ*_d_ between droplets’ velocity angles and their azimuth angles relative to the aggregate. (F) Plot of maximum flow velocities within droplet condensates as a function of droplet-to-aggregate center distance *d*. (G) Plot of maximum flow velocities within droplet condensates as a function of the center distance *d* divided by the droplet radius *r, d*/*r*.

Our theoretical analysis further suggests that these active flows originate from Marangoni effects driven by interfacial tension gradients at droplet-aggregate interfaces (details in the supplementary materials). The theoretical model suggests that the maximum internal velocity *V*_*c*_ is proportional to the Marangoni number (Ma), which scales with the ratio of droplet radius *r* to droplet-aggregate center distance *d* when *d* ≫ *r*, as *V*_*c*_ ∼ Ma ∼ *r*/*d* (details in the supplementary materials). Consequently, replotting *V*_*c*_ against *d*/*r* collapses data from three independent experiments onto a theoretical curve (Fig. 4G). This good agreement between experiments and theory underscores the critical role of Marangoni effects in causing active flows and driving long-range chemotaxis ripening of ATP condensates.

### Dialytaxis of droplets into aggregates

Not only does the Marangoni effect drive internal fluid flow within sessile droplet condensates, it can also cause these droplets to slide when the surface is more hydrophobic and consequently offers lower friction (Fig. 5A). On hydrophobic slides treated with fluorosilane coatings, condensate droplets migrate gradually toward and eventually merge with the central aggregate (Fig. 5B, Fig. 5C, and movie S8). In particular, droplet velocities increase as they approach the fibril cluster (Fig. 5D). As a result, the mobility of droplets accelerates aggregate growth slightly, with its projected area increasing over time with a power-law exponent greater than 1 (Fig. S15). In the later stage, however, the growth slows down, primarily due to the depletion of surrounding droplets (Fig. S15). The motion of condensates down dilute-phase concentration gradients has been recently observed for externally imposed concentration gradients, where condensates swim towards conditions that favor their dissolution [32]. This directed motion has been proposed to be caused by the Marangoni effect, which is known to play a causative role in droplet swimming [33].

**Fig 5.**
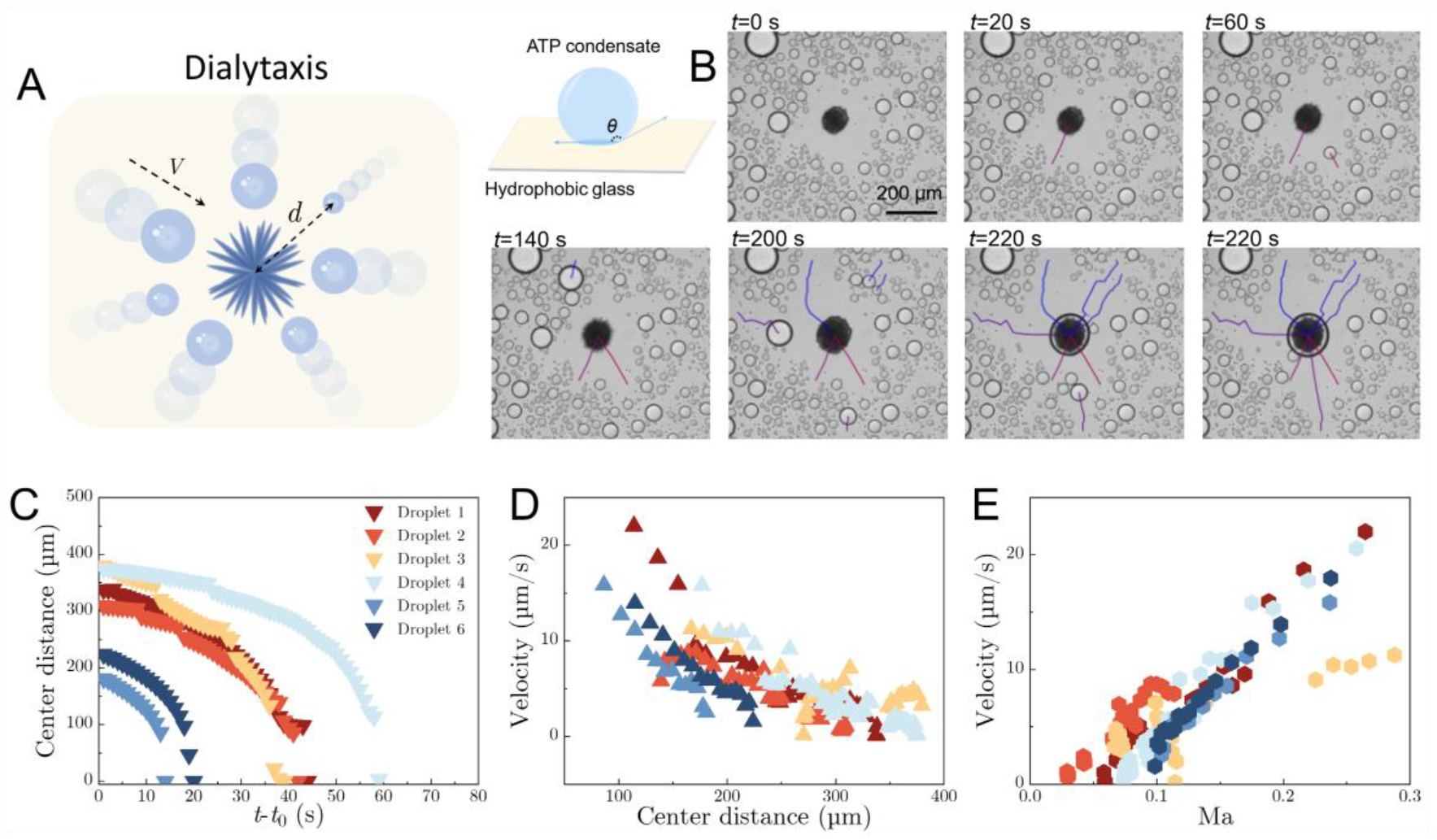
Dialytaxis of droplets into aggregates. (A) Schematic of condensates sliding toward a central aggregate on a hydrophobic surface. (B) Bright-field images and trajectory overlays show collective dialytaxis motions of neighboring droplets toward a central aggregate along chemotaxis gradients. (C) Plot of droplet-to-aggregate center distance as function of time for six independent droplets. (C) Plot of droplet velocities versus the droplet-to-aggregate center distance. (E) Plot of droplet velocities against Marangoni number *Ma*.

Remarkably, the concentration gradient in our study is endogenously generated by the diffusion-limited growth of the fibril aggregate. The dilute phase concentration decreases approaching the aggregates, and the concentration gradient magnitude increases (details in the supplementary materials). To validate this, we have measured the droplet radius *r* and center-to-center distance *d* from the nearest aggregate over time for several droplets in Fig. 5B. From these acquired data we are again able to calculate the Marangoni number (Ma) over time, where Ma 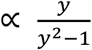 with 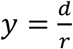 (details in the supplementary materials). Plotting droplet velocities against Ma over time, we found data from different droplets largely collapse onto a straight line (Fig. 5E), though some outliers occur when droplets become stuck behind other droplets or when droplets merge with one another or the aggregate at the end (Fig. S16 and Movie S8). This confirms the motion of these droplets shares same driving force as explored in [32] and as established for the internal active flow of aging condensates in this work.

## Discussion

This study demonstrates that aging condensates exhibit a remarkably rich and complex repertoire of dynamic behaviors, including wetting-induced leaps, chemotaxis ripening, active internal flows, and dialytaxis motion. These new aging mechanisms accelerate the condensate aging timescale from hours-to-days reported previously to minutes in this study [14, 18]. The chemotaxis ripening during condensate aging is reminiscent of Ostwald ripening, a canonical process in inhomogeneous droplet suspensions (e.g., water-in-oil emulsions) [34], although this mechanism has been suggested to be negligible for condensates [35, 36]. The Ostwald ripening is driven by Laplace pressure 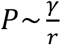 between droplets of unequal sizes [34, 37], thereby suggesting an inverse size dependence, where smaller droplets dissolve preferentially and transfer their components to larger neighbors. Unlike the inverse size dependence in Ostwald ripening, the chemotaxis ripening effect studies here exhibits a positive correlation with droplet radius: larger droplets experience stronger Marangoni effects and exhibit more vigorous internal active flows. This chemotaxis ripening is driven by non-equilibrium physicochemical gradients, such as the difference in ATP chemical potentials and interfacial tension gradients between droplets and aggregates.

These dynamic behaviors observed in this study may represent universal features of condensate aging but are potentially masked in condensate systems with sluggish aging dynamics. For example, FUS protein condensates, implicated in amyotrophic lateral sclerosis (ALS) disease, undergo phase transitions to form similar fibrous structures over hours [14, 21]. In addition, the interaction between fibrillar structures and condensates could drive their structural remodeling [38-41], a process that mirrors the wetting-induced colocalization of droplets and aggregates observed here. This study also challenges the classic view of condensates as passive entities; instead, aging condensates act as active droplets, sensing chemical gradients and responding by directed migration via chemotaxis and dialytaxis mechanisms, even under global equilibrium conditions. Overall, these findings provide crucial mechanistic insights into condensate aging with implications for understanding their roles in the pathogenesis of neurodegenerative diseases.

## Competing interest

Ho Cheung Shum is a scientific advisor of EN Technology Limited, MicroDiagnostics Limited, PharmaEase Tech Limited, Upgrade Biopolymers Limited and Multera Limited, in which he owns some equity, and is a founding director and co-director of the research center, Advanced Biomedical Instrumentation Centre Limited. The works in this paper are, however, not directly related to the works of these entities, as far as we know. The authors declare no other competing interests.

## Acknowledgements

We thank Tuomas Knowles and David Weitz for helpful discussions and comments on this project. This work was supported by the General Research Fund (Nos. 17306221, and 17317322), and Collaborative Research Fund (C7165-20GF) from the Research Grants Council (RGC) of Hong Kong, as well as the National Natural Science Foundation of China (NSFC)-RGC Joint Research Scheme (N_HKU718/19), ETH Zurich (AD, TCTM) and the Swiss National Science Foundation (grant no SNF 219703 to TCTM). H.C.S. was funded in part by the RGC Senior Research Fellow (SRFS2425-7S04) by RGC and the Health@InnoHK program of the Innovation and Technology Commission of the Hong Kong SAR Government.

## Author contributions

F.C. conceived and designed the research; F.C., P.D.W., and Y.W. performed the experiments and analyzed data; A.J.D. and T.C.T.M developed theoretical models. F.C., T.C.T.M, and H.C.S. supervised the research.

## Data Availability

All data needed to evaluate the conclusions in the paper are present in the paper and/or the Supplementary Information, and also from the corresponding author upon request.

## References

1. Cho, W.-K., et al., Mediator and RNA polymerase II clusters associate in transcription-dependent condensates. Science, 2018. 361(6400): p. 412–415.

2. Hnisz, D., et al., A phase separation model for transcriptional control. Cell, 2017. 169(1): p. 13–23.

3. Zeng, M., et al., Reconstituted postsynaptic density as a molecular platform for understanding synapse formation and plasticity. Cell, 2018. 174(5): p. 1172-1187. e16.

4. Wu, X., et al., Liquid-liquid phase separation in neuronal development and synaptic signaling. Developmental cell, 2020. 55(1): p. 18–29.

5. Klosin, A., et al., Phase separation provides a mechanism to reduce noise in cells. Science, 2020. 367(6476): p. 464–468.

6. Chen, F., et al., Size Scaling of Condensates in Multicomponent Phase Separation. Journal of the American Chemical Society, 2024.

7. Boyd-Shiwarski, C.R., et al., WNK kinases sense molecular crowding and rescue cell volume via phase separation. Cell, 2022. 185(24): p. 4488-4506.e20.

8. Pak, C.W., et al., Sequence determinants of intracellular phase separation by complex coacervation of a disordered protein. Molecular cell, 2016. 63(1): p. 72–85.

9. Elbaum-Garfinkle, S., et al., The disordered P granule protein LAF-1 drives phase separation into droplets with tunable viscosity and dynamics. Proceedings of the National Academy of Sciences, 2015. 112(23): p. 7189–7194.

10. Qamar, S., et al., FUS phase separation is modulated by a molecular chaperone and methylation of arginine cation-π interactions. Cell, 2018. 173(3): p. 720-734. e15.

11. Martin, E.W., et al., Valence and patterning of aromatic residues determine the phase behavior of prion-like domains. Science, 2020. 367(6478): p. 694–699.

12. Brangwynne, C.P., et al., Germline P granules are liquid droplets that localize by controlled dissolution/condensation. Science, 2009. 324(5935): p. 1729–1732.

13. Lyon, A.S., W.B. Peeples, and M.K. Rosen, A framework for understanding the functions of biomolecular condensates across scales. Nature Reviews Molecular Cell Biology, 2021. 22(3): p. 215–235.

14. Patel, A., et al., A liquid-to-solid phase transition of the ALS protein FUS accelerated by disease mutation. Cell, 2015. 162(5): p. 1066–1077.

15. Lin, Y., et al., Formation and maturation of phase-separated liquid droplets by RNA-binding proteins. Molecular cell, 2015. 60(2): p. 208–219.

16. Alberti, S. and A.A. Hyman, Biomolecular condensates at the nexus of cellular stress, protein aggregation disease and ageing. Nature reviews Molecular cell biology, 2021. 22(3): p. 196–213.

17. Boija, A., I.A. Klein, and R.A. Young, Biomolecular condensates and cancer. Cancer cell, 2021. 39(2): p. 174–192.

18. Jawerth, L., et al., Protein condensates as aging Maxwell fluids. Science, 2020. 370(6522): p. 1317–1323.

19. Alshareedah, I., et al., Sequence-specific interactions determine viscoelasticity and ageing dynamics of protein condensates. Nature Physics, 2024. 20(9): p. 1482–1491.

20. Zhang, H., et al., RNA controls PolyQ protein phase transitions. Molecular cell, 2015. 60(2): p. 220–230.

21. Boczek, E.E., et al., HspB8 prevents aberrant phase transitions of FUS by chaperoning its folded RNA-binding domain. Elife, 2021. 10: p. e69377.

22. Abbas, M., et al., A short peptide synthon for liquid–liquid phase separation. Nature Chemistry, 2021: p. 1–9.

23. Yuan, C., et al., Nucleation and growth of amino acid and peptide supramolecular polymers through liquid–liquid phase separation. Angewandte, 2019. 131(50): p. 18284–18291.

24. Ratke, L. and P.W. Voorhees, Growth and coarsening: Ostwald ripening in material processing. 2002: Springer Science & Business Media.

25. Krainer, G., et al., Reentrant liquid condensate phase of proteins is stabilized by hydrophobic and non-ionic interactions. Nature Communications, 2021. 12(1): p. 1–14.

26. Smokers, I.B., et al., Selective ion binding and uptake shape the microenvironment of biomolecular condensates. Journal of the American Chemical Society, 2024.

27. Folkmann, A.W., et al., Regulation of biomolecular condensates by interfacial protein clusters. Science, 2021. 373(6560): p. 1218–1224.

28. Feric, M., et al., Coexisting liquid phases underlie nucleolar subcompartments. Cell, 2016. 165(7): p. 1686–1697.

29. Michaels, T.C.T., et al., Dynamics of oligomer populations formed during the aggregation of Alzheimer’s Aβ42 peptide. Nature Chemistry, 2020. 12(5): p. 445–451.

30. Zwicker, D., et al., Growth and division of active droplets provides a model for protocells. Nature Physics, 2017. 13(4): p. 408–413.

31. Adkins, R., et al., Dynamics of active liquid interfaces. Science, 2022. 377(6607): p. 768–772.

32. Jambon-Puillet, E., et al., Phase-separated droplets swim to their dissolution. Nature communications, 2024. 15(1): p. 3919.

33. Schmitt, M. and H. Stark, Marangoni flow at droplet interfaces: Three-dimensional solution and applications. Physics of Fluids, 2016. 28(1).

34. Yao, J.H., et al., Theory and simulation of Ostwald ripening. Physical review B, 1993. 47(21): p. 14110.

35. Nakashima, K.K., et al., Active coacervate droplets are protocells that grow and resist Ostwald ripening. Nature Communications, 2021. 12(1): p. 1–11.

36. Lee, D.S., N.S. Wingreen, and C.P. Brangwynne, Chromatin mechanics dictates subdiffusion and coarsening dynamics of embedded condensates. Nature Physics, 2021. 17(4): p. 531–538.

37. Zhang, Y., et al., Mechanical frustration of phase separation in the cell nucleus by chromatin. Physical review letters, 2021. 126(25): p. 258102.

38. Pombo-García, K., et al., Membrane prewetting by condensates promotes tight-junction belt formation. Nature, 2024. 632(8025): p. 647–655.

39. Mangiarotti, A., et al., Wetting and complex remodeling of membranes by biomolecular condensates. Nature Communications, 2023. 14(1): p. 2809.

40. Setru, S.U., et al., A hydrodynamic instability drives protein droplet formation on microtubules to nucleate branches. Nature physics, 2021. 17(4): p. 493–498.

41. Gouveia, B., et al., Capillary forces generated by biomolecular condensates. Nature, 2022. 609(7926): p. 255–264.

