## Supplementary information for "Endogenous flow and dialytaxis govern aging of Adenosine 5’-triphosphate (ATP) condensates"

### I. MATERIALS AND METHODS

**Chemicals:** Adenosine 5′-triphosphate (ATP, A26209), Polyethylene glycol (PEG) (Mw ~8000, 89510), ethyl alcohol (ethanol, 459844), Fluorescein isothiocyanate–dextran (FITC-dextran, Mw ~10, 000, FD10S), Calcium chloride (C4901), Poly-L-lysine–FITC Labeled (FITC-PLL, Mw ~30,000–70,000, P3069), Rhodamine 6G (R4127) were purchased from Sigma-Aldrich. Calcein (C794029) and Nile red (N815046) were purchased from Macklin. Rhodamine-labeled PEG (Rh-PEG, Mw ~10,000, R164224) were purchased from Aladdin. Sodium Chloride (5M, AM9759), potassium chloride (2M, AM9640G), Magnesium chloride (1 M, AM9530G), UltraPure Water (10977015) were purchased from ThermoFisher Scientific. Fluorescent polystyrene microspheres (diameter: 1 μm, 10mg/ml, excitation peak: 488 nm, 7-31-0100) were purchased from Tianjin Junvijia Technology Co. Ltd. Glass slides with fluorosilane coating were purchased from Cytonix. UltraPure Water was used in all experiments. All chemicals were used as received without further purification.

**Preparation of aging condensates**: ATP was dissolved in water to prepare a 500 mM stock solution. The ATP solution was aliquoted, stored at −20 °C, and kept on ice during experimental use. PEG 8000 was dissolved in water to prepare a 500 mg/mL stock solution. To generate droplet condensates, water, ATP, PEG and ethanol were sequentially added into a microcentrifuge tube according to specific experimental conditions to a final volume of 40 μL. For PIV experiments, 1 μL solution of fluorescent microparticles [xxx mg/ml] was added to the mixture. For partitioning experiments, 1 μL fluorophore solution was added to the mixture. After each addition, the mixture was gently pipette-mixed and vortexed (~2000 rpm, ~10 s).

**Imaging and characterization:** The optical images of condensates were captured using an inverted Leica microscope. Three-dimensional images of ATP aggregates were captured by a confocal laser scanning microscope (CLSM, Carl Zesis LSM 700). Coverslides were first cleaned with ethanol and water to remove contaminants before use. ~8 μL condensate solution was added into a closed chamber that was customized using a spacer sandwiched by two coverslides at a 120 μm gap. The sample within the chamber was then observed under optical or fluorescence microscopy to monitor the aging dynamics of condensates. The surface tension of ATP aqueous solutions was measured experimentally by the pendant drop method using a customized MATLAB code.

**Condensate Growth Dynamics**: Videos recorded at 15 frames per second were cropped, trimmed and exported as individual frames before being analyzed with MATLAB. Frames were binarized, noise-filtered, and spatially calibrated using pixel-to-mm conversion (scale ratio: k = μm/pixel) to segment condensates and aggregates. The area and center position of aggregates were determined using their boundary data, while for condensates, circles were detected. This, combined with temporal resolution using frame rate, enabled area, velocity, and center position calculations over time.

**Particle Image Velocimetry:** Videos recorded at 15 frames per second were cropped, trimmed and exported as individual frames before analyzing with Particle Image Velocimetry (MATLAB). Cropped video sequences were imported, and every $n^{th}$ frame was processed to capture flow evolution. Spatial calibration was performed by drawing a reference line across the image width, converting pixel coordinates to physical dimensions (μm/pixel) using the original video scale. A mask was optionally applied to exclude static regions or artifacts. Contrast settings were optimized via histogram equalization and intensity thresholding to enhance tracer particle visibility. Multi-pass cross-correlation analysis employed interrogation areas progressively halved across passes with FFT window deformation to refine displacement accuracy. Velocity magnitudes ($m \cdot s^{-1}$) were scaled using spatial calibration data, with color maps adjusted for ranges relevant to condensate flows. Vector arrow density and scaling were optimized to prevent overlapping while maintaining directional clarity.

**Single particle tracking:** Images from recorded videos (typically recorded at 15 frames per second) were exported and preprocessed using MATLAB. Single particle tracking was performed, with the x-y coordinates of each microparticle extracted using custom scripts to obtain particle trajectories. The mean squared displacement (MSD) for each trajectory was calculated using the formula:

$$\mathrm{MSD}(\Delta t) = \langle (x(t + \Delta t) - x(t))^2 \rangle + \langle (y(t + \Delta t) - y(t))^2 \rangle.$$

where $x$ and $y$ denote the centroid coordinates of the particle and $\Delta t$ indicates the lag time. MSD results are generated for individual particles within a single condensate and averaged to minimize errors. Particles around the center of droplets were selected and analyzed to avoid interface artifacts. To determine the 2D diffusion coefficient (D), the MSD curve was fitted with $\mathrm{MSD}(\Delta t) = 4D\Delta t$. The viscosity within condensates can be determined using the Stokes–Einstein relation.

### II. DIFFUSION-LIMITED FIBRIL CLUSTER GROWTH KINETICS

We describe diffusion-limited growth of a fibril cluster with an approximately spherical shape. We assume that fibril growth is diffusion-limited (after an initial transient dynamics time) and that the fibrils occupy a roughly spherical space and are oriented radially. As a consequence, the fibril cluster can be modelled as an absorbing sphere with radius $R$, such that as soon as a monomer reaches the cluster, it is incorporated in it. The steady-state concentration profile $c(\boldsymbol{x})$ of monomers outside of the fibrils is governed by the diffusion equation

$$D\nabla^2 c(x) = 0\,, \tag{1}$$

Where $D$ is the diffusion coefficient of monomers in solution. Using spherical coordinates, the equation becomes

$$\frac{D}{r^2}\frac{\partial}{\partial r}\left(r^2\frac{\partial c(r)}{\partial r}\right) = 0\,, \tag{2}$$

where $r$ is the radial distance from the fibril cluster. We impose the following boundary conditions

$$c(r = \infty) = c_\infty \tag{3a}$$

$$c(r = R) = 0\,, \tag{3b}$$

where $c_\infty$ is the bulk monomer concentration. The solution to Eq. (2) is

$$c(r) = c_\infty\left(1 - \frac{R}{r}\right). \tag{4}$$

Now, the flux at the fibril cluster boundary, $\mathrm{r} = \mathrm{R}$, is

$$J(r = R) = D\left.\frac{\partial c(r)}{\partial r}\right|_{r=R} = \frac{D\,c_\infty}{R}\,. \tag{5}$$

Integrated over the cluster surface the total flux (number of monomers per unit time) is then:

$$\frac{dN}{dt} = 4\pi D R c_\infty, \tag{6}$$

where $N$ is the concentration of protein in fibrils. Assuming a spherical fibril cluster, $N$ is proportional to the volume of the sphere, i.e.:

$$N\,\nu_0 = \frac{4}{3}\pi R^3, \tag{7}$$

where $\nu_0$ is the volume of one monomer. We therefore have:

$$R\,\frac{dR}{dt} = D\,c_\infty\,\nu_0 \tag{8}$$

with solution

$$R(t)^2 - R(0)^2 = 2D\,c_\infty\,\nu_0\,t\,, \tag{9}$$

which is the famous $R^2$-law for diffusion-limited growth. The area of the fibril cluster therefore grows in proportion to $t$.

### III. ACTIVE FLOWS AND DIALYTAXIS OF AGING CONDENSATES

We anticipate in the main text that both internal flow within droplets and droplet motion toward fibril clusters are driven by the Marangoni effect. Marangoni flow is, in turn, driven by surface tension gradients. We expect such a gradient to exist here as a consequence of the concentration gradient outside the droplet, which itself arises from the diffusion-limited growth of the fibril cluster.

The first question is how does surface tension $\gamma$ depend on the concentration outside the interfacial region in the dilute phase? We assume (very reasonably) that the concentration inside the droplet remains constant at roughly its phase equilibrium value $c^I$. Crudely, the surface tension should scale with the drop in concentration across the interface. Thus $\gamma \approx A_1(c^I - c^{II})$.

The magnitude of the Marangoni flow is governed by the Marangoni number Ma:

$$\mathrm{Ma} = \left|\frac{d\gamma}{dX}\right| \frac{\Delta X L}{\mu D_X}, \tag{10}$$

where $X$ is the quantity whose gradient induces the interfacial shear, in this case $c^{II}$, $D_X$ is its diffusion constant, and $\mu$ is the viscosity. $d\gamma/dc^{II}$ is easily evaluated and this formula thus reduces to:

$$\mathrm{Ma} = A_1 \frac{\Delta c^{II} L}{\mu D}. \tag{11}$$

The remaining quantities to evaluate are $L$, the length scale of the gradient of the quantity ($c^{II}$) inducing interfacial shear, and $\Delta c^{II}$ is the total change in $c^{II}$ across the droplet.

We assume the flux out of the droplet is slow relative to the reaction-diffusion process in the dilute phase that governs fibril cluster growth. Based on the data this assumption is very reasonable, as otherwise the droplets would not be stable for all but the shortest timescales near the fibrils. Then $c^{II}$ is given by the equation above, Eq. (4):

$$c^{II}(r) = c_\infty\left(1 - \frac{R}{r}\right). \tag{12}$$

So, $\Delta c^{II}$ across the droplet is given by

$$\Delta c^{II} = c_\infty \left(1 - \frac{R}{r+d}\right) - c_\infty \left(1 - \frac{R}{r-d}\right), \tag{13}$$

where $r$ is the distance between the centers of the droplet and the fibril cluster, $d$ is the droplet radius, and $R$ is the fibril cluster radius. Simplified, this becomes:

$$\Delta c^{II} = c_\infty R \left(\frac{1}{r-d} - \frac{1}{r+d}\right) = \frac{2c_\infty R\, d}{r^2 - d^2}. \tag{14}$$

Finally, the characteristic length scale of the concentration gradient is $L = r$. Putting everything together we get:

$$\text{Ma} \propto \frac{y}{y^2 - 1}, \text{ where } y = \frac{r}{d}. \tag{15}$$

Since generally $y \gg 1$, this predicts a $1/r$ scaling of the flow velocity inside the droplet. It also predicts a $1/r$ scaling of the droplet velocity towards the fibril cluster when droplets are on hydrophobic surfaces. Both of these predictions are consistent with our experimental observations (see main text).

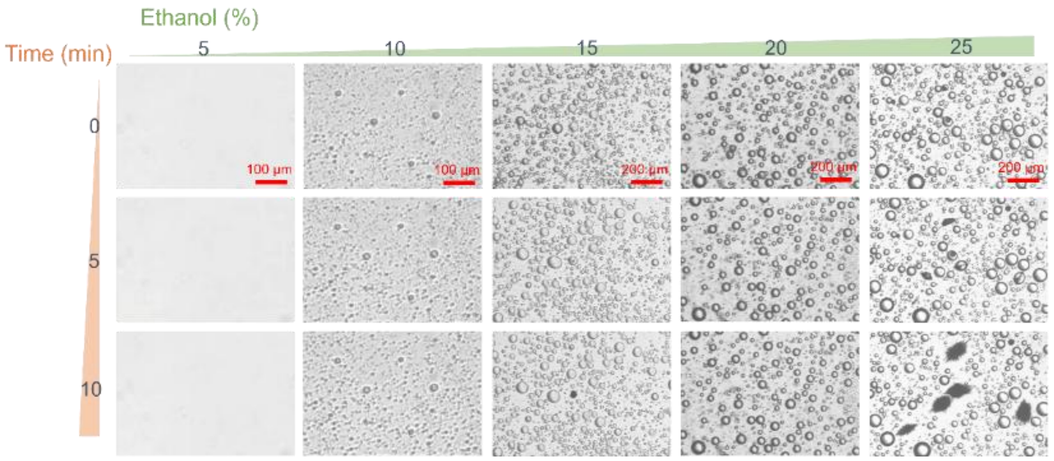


**Fig. S1.** The effect of ethanol volume fraction on the aging of ATP condensates. Bright-field images of mixtures of ATP (100 mM), PEG (200mg/ml), and different volume fractions of ethanol, from 5% v/v to 25% v/v. More droplets and aging ATP condensates are observed with increasing volume fraction of ethanol.

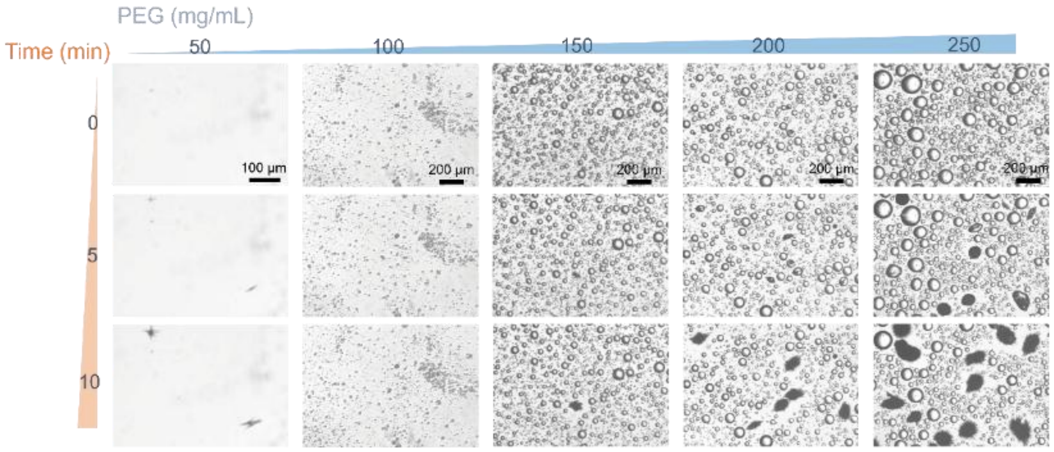


**Fig. S2.** The effect of PEG concentration on the aging of ATP condensates. Bright-field images of mixtures of ATP (100 mM), ethanol (25% v/v), and different concentrations of PEG (mg/ml). More droplets and aging ATP condensates are observed with increasing volume fraction of ethanol.

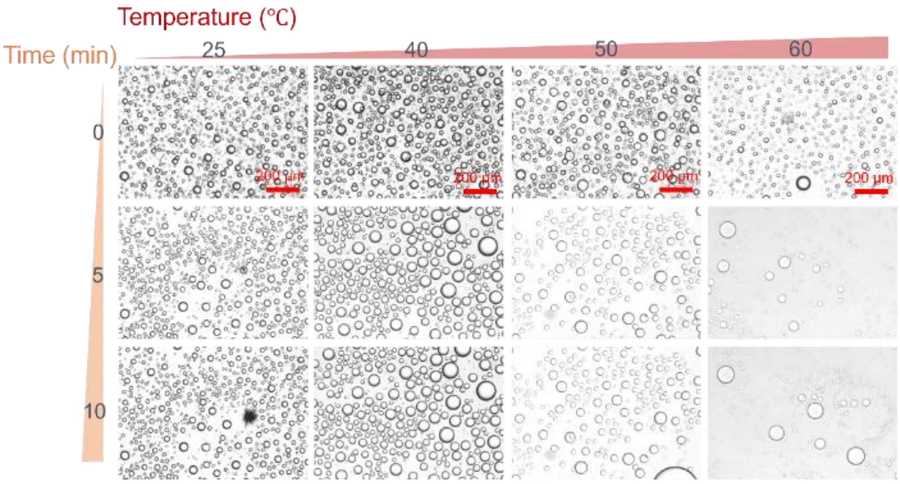


**Fig. S3.** The effect of temperature on the aging of ATP condensates. Condensates are formed at: ATP (100 mM), PEG (200 mg/ml), and ethanol (25% v/v). Aging becomes suppressed at high temperatures.

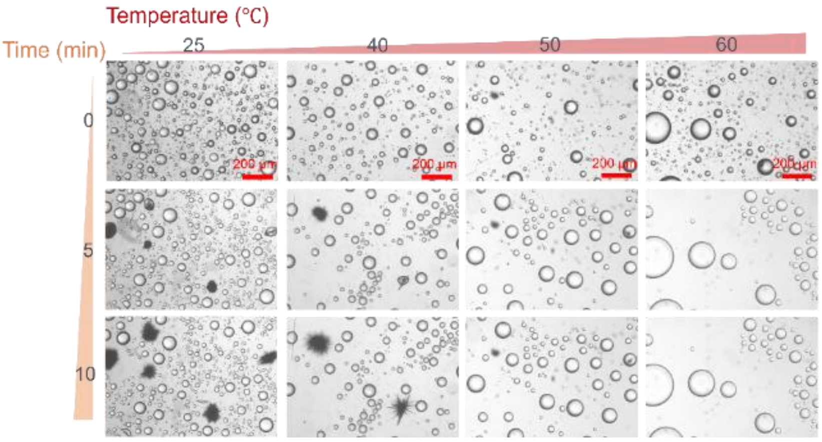


**Fig. S4.** The effect of temperature on the aging of ATP condensates. Condensates are formed at ATP (100 mM), PEG (250 mg/ml), and ethanol (25% v/v). Aging becomes suppressed at high temperatures.

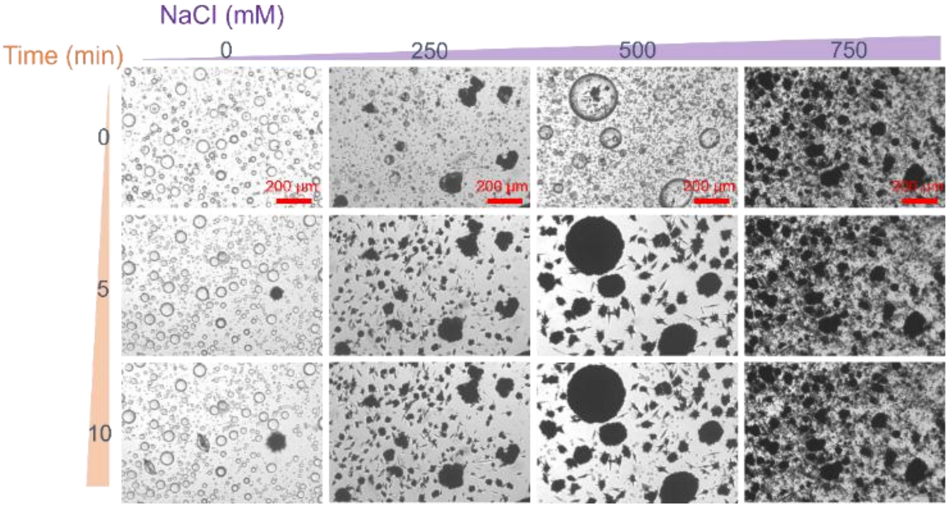


**Fig. S5.** The effect of monovalent sodium chloride (NaCI) on the aging of ATP condensates. Condensates are formed at: ATP (100 mM), PEG (200 mg/ml), and ethanol (25% v/v).

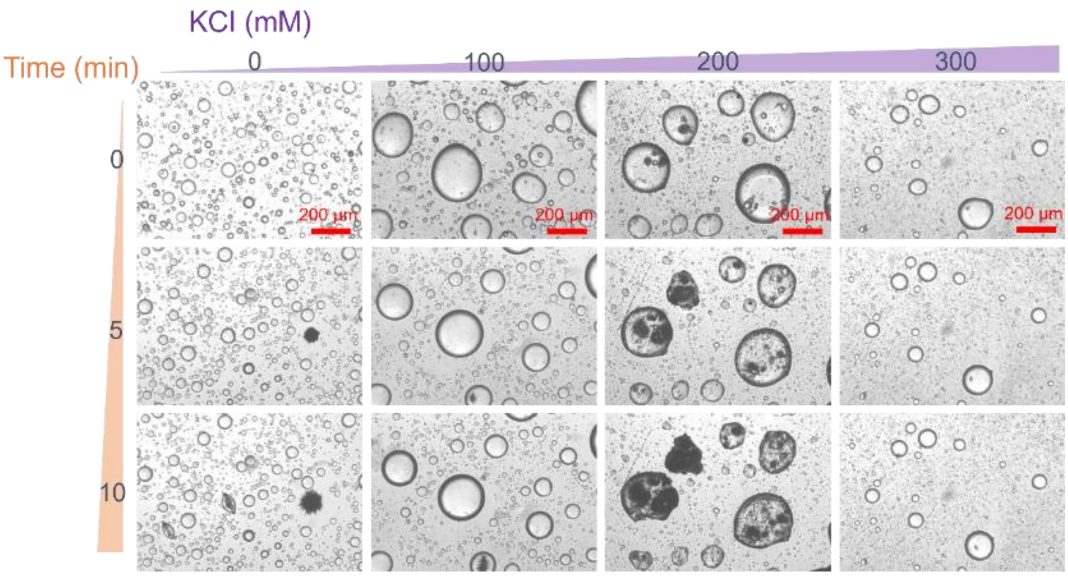


**Fig. S6.** The effect of monovalent potassium chloride (KCl) on the aging of ATP condensates. Condensates are formed at: ATP (100 mM), PEG (200 mg/ml), and ethanol (25% v/v).

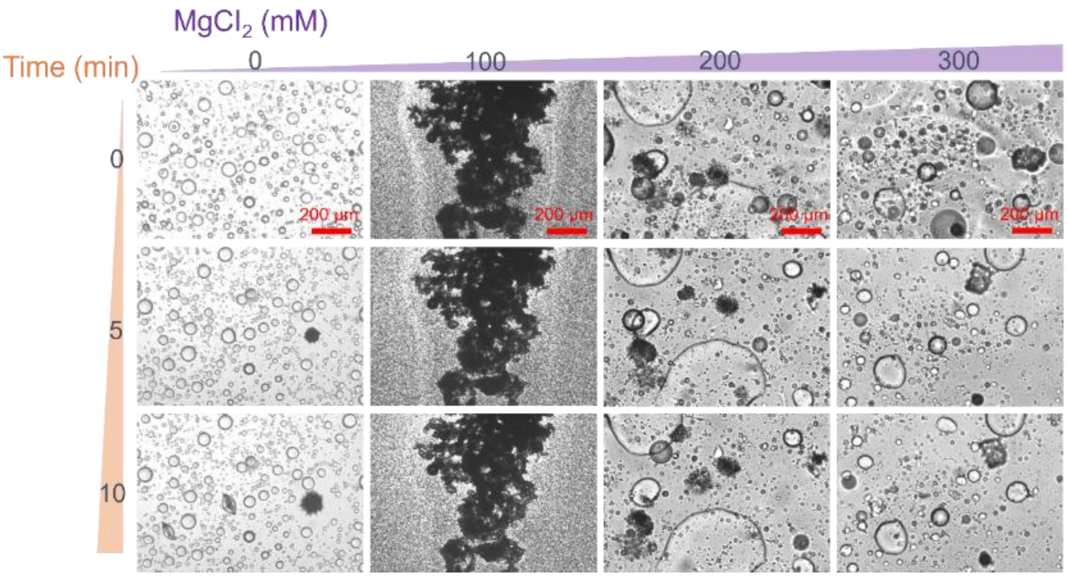


**Fig. S7.** The effect of divalent Magnesium chloride ($MgCI_2$) on the aging of ATP condensates. Condensates are formed at: ATP (100 mM), PEG (200 mg/ml), and ethanol (25% v/v).

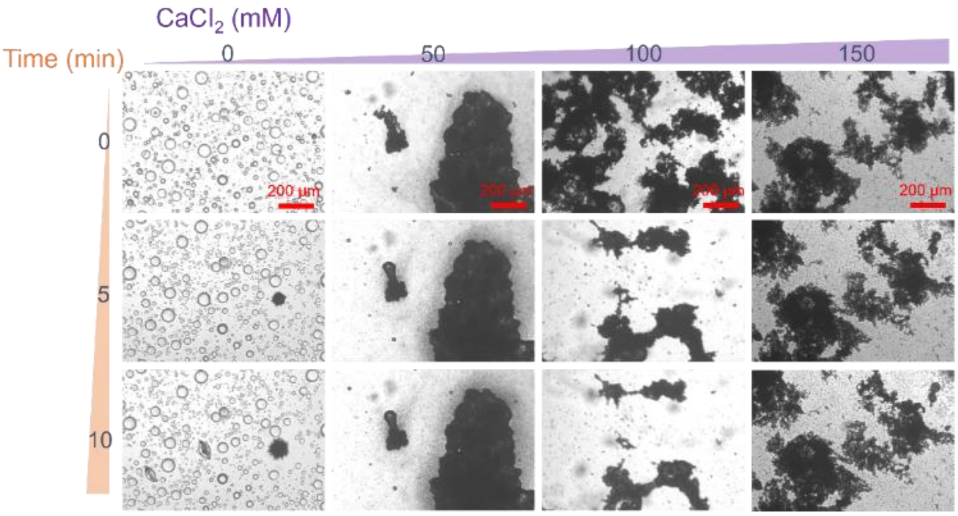


**Fig. S8.** The effect of monovalent calcium chloride (CaCI2) on the aging of ATP condensates. Condensates are formed at: ATP (100 mM), PEG (200 mg/ml), and ethanol (25% v/v).

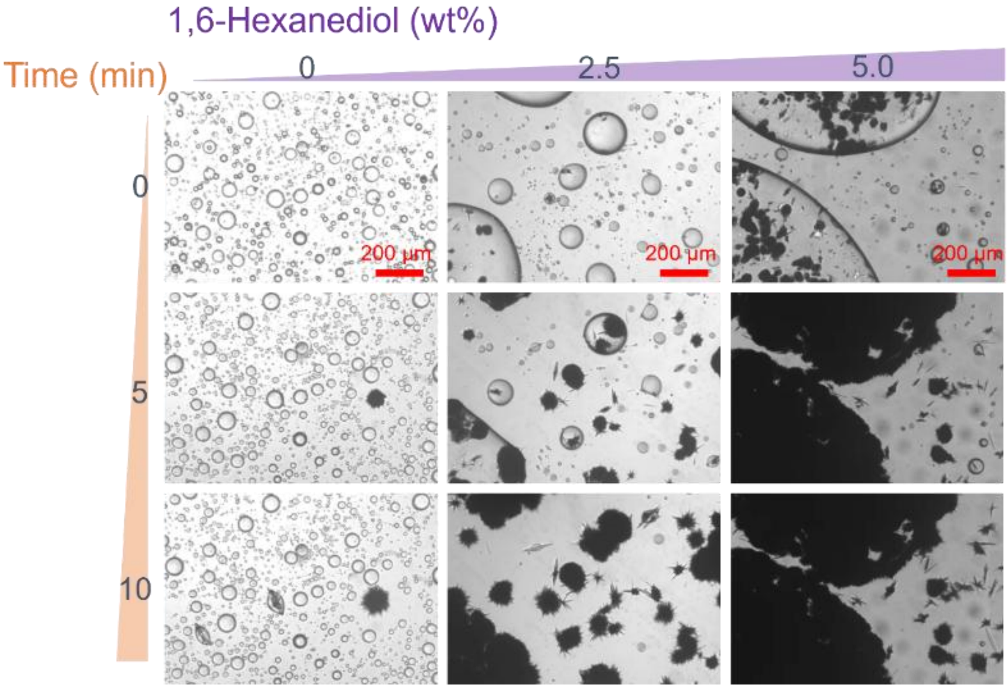


**Fig. S9.** The effect of 1,6 Hexanediol on the aging of ATP condensates. Condensates are formed at: ATP (100 mM), PEG (200 mg/ml), and ethanol (25% v/v).

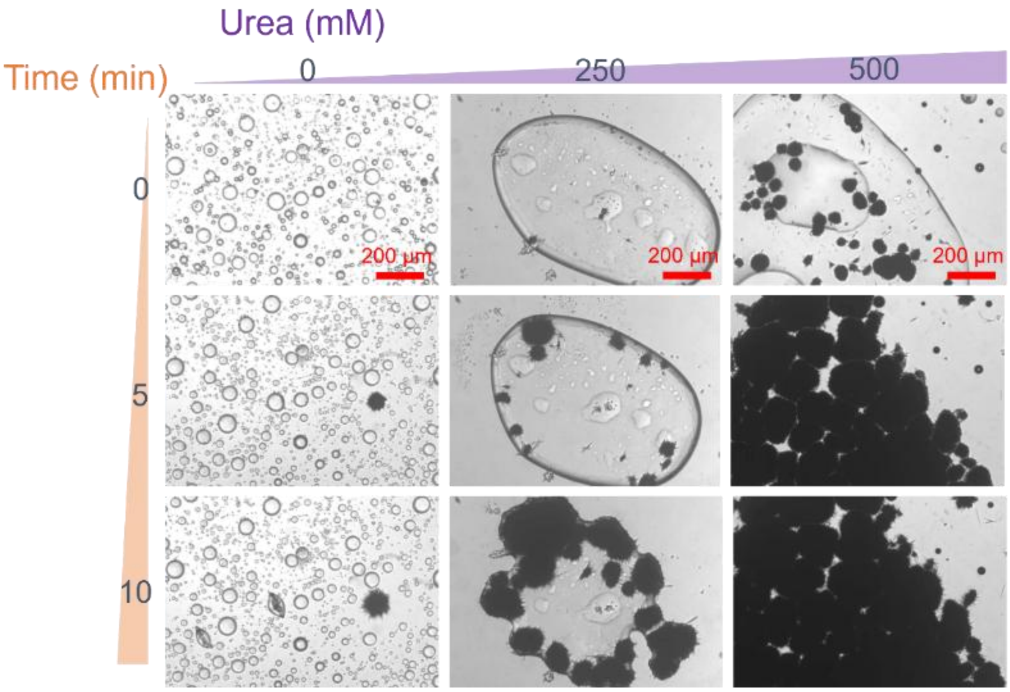


**Fig. S10.** The effect of Urea on the aging of ATP condensates. Condensates are formed at: ATP (100 mM), PEG (200 mg/ml), and ethanol (25% v/v).

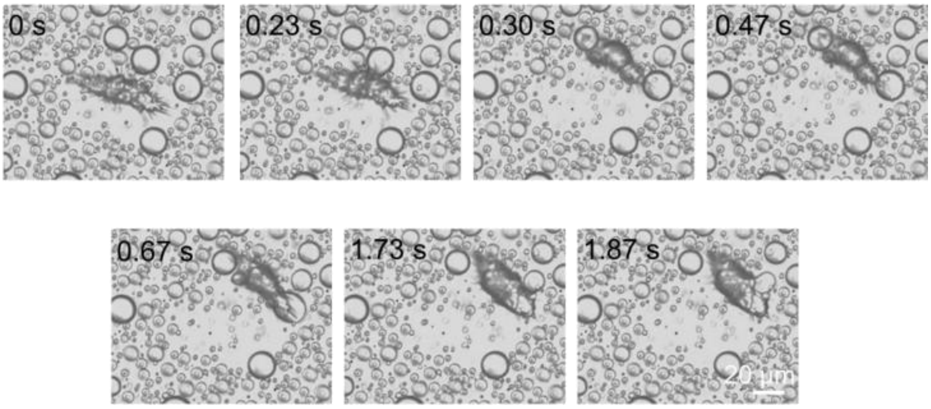


**Fig. S11.** Bright-field images showing the leap and colocalization of an aggregate with neighboring droplets by direct contact. These images correspond to movie S3.

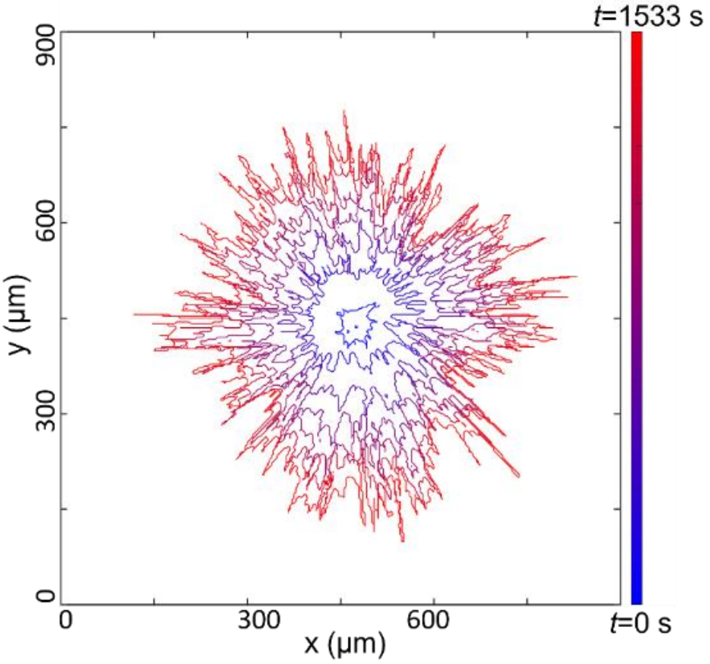


**Fig. S12.** The boundary profiles of aggregates over time were extracted using customized codes.

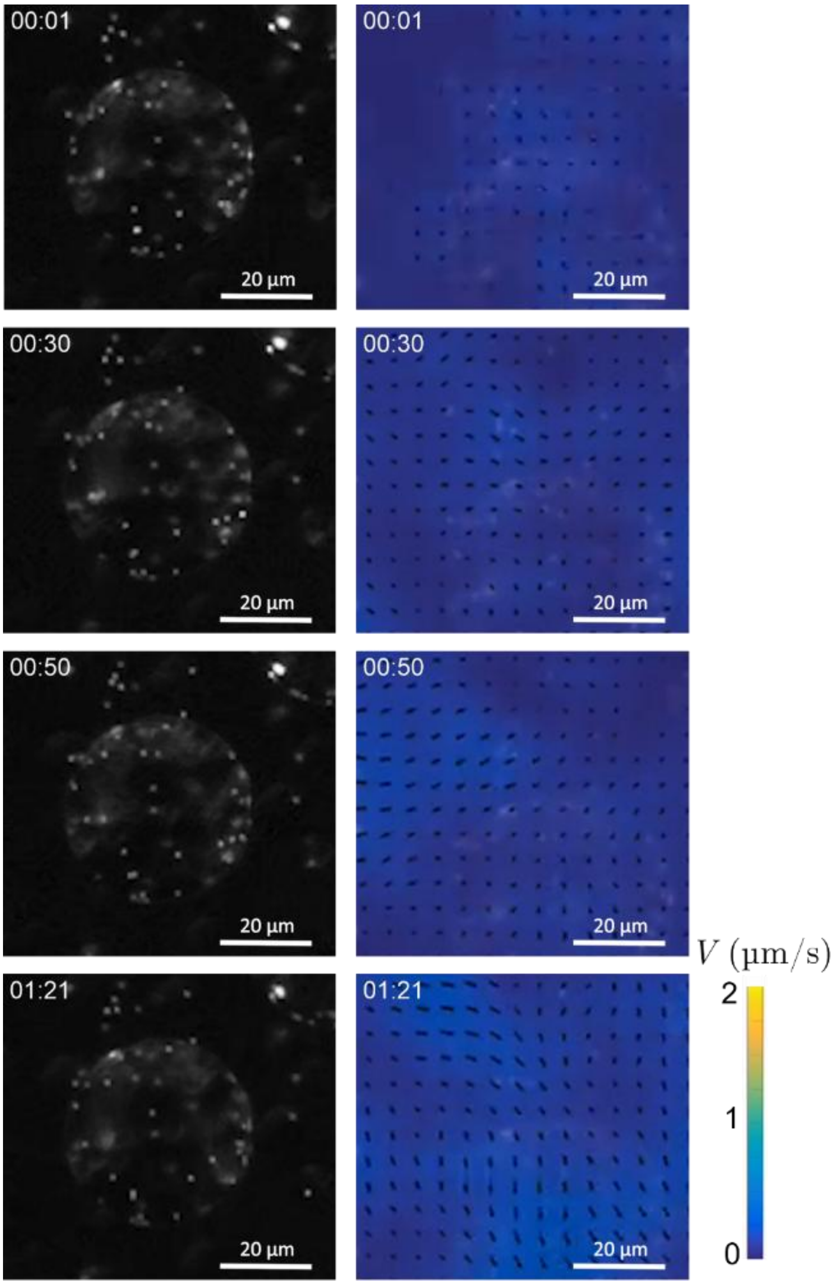


**Fig. S13.** PIV image of flow fields within a non-aging condensate. Black arrows denote the velocity magnitude and direction of flow fields.

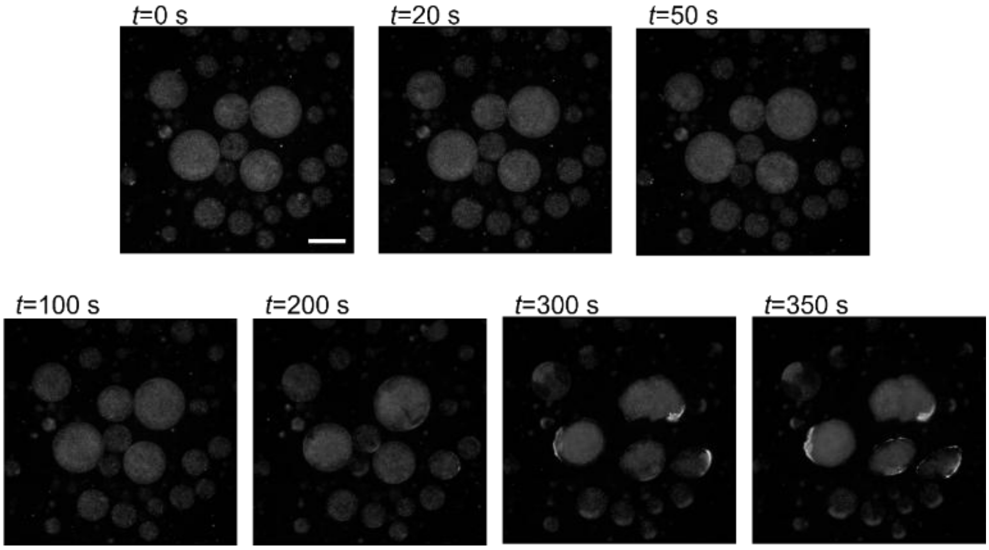


**Fig. S14.** Fluorescence images show that microparticles become immobile when ATP droplets transition into aggregates. The scale bar is 20 μm.

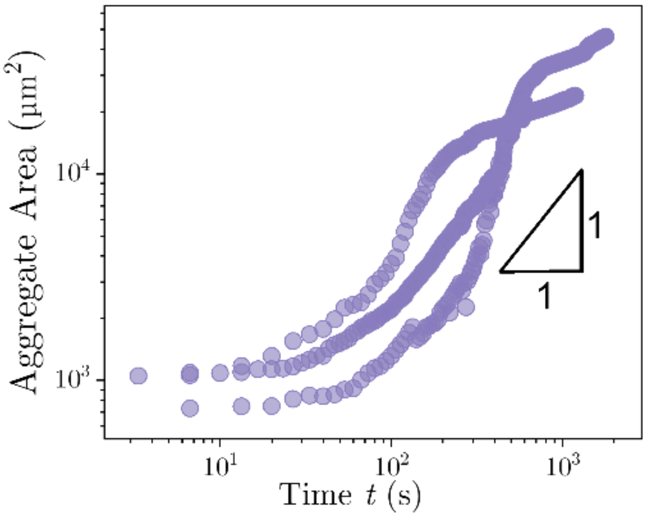


**Fig. S15.** Time-evolution of the area of aggregate that forms on a hydrophobic substrate in which condensates undergo dialytaxis into aggregates in Fig. 2H.

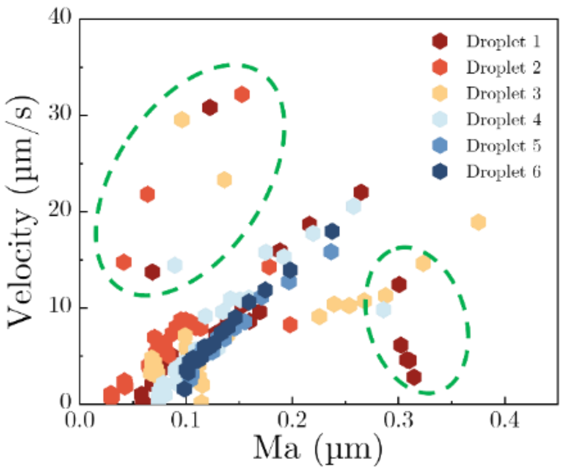


**Fig. S16.** (A) Plot of droplet velocities against Marangoni number *Ma* for six independent droplets undergoing dialytaxis motions on a hydrophobic surface. Outliers marked in green circles occur when droplets become stuck behind other droplets or when droplets merge with one another or with the aggregate at the end.
